# Matrix metalloproteinases and tissue inhibitors of metalloproteinases in murine coronavirus-induced neuroinflammation

**DOI:** 10.1101/2020.09.17.302877

**Authors:** Sourodip Sengupta, Sankar Addya, Diptomit Biswas, Jayasri Das Sarma

## Abstract

Mouse hepatitis virus (MHV) belongs to the same beta-coronavirus family as SARS-CoV-2, MERS-CoV, and SARS-CoV. Studies have shown the requirement of host cellular proteases for priming the surface spike protein during viral entry and transmission in coronaviruses. The metzincin family of metal-dependent endopeptidases called matrix metalloproteinases (MMPs) is involved in virus encephalitis, enhanced blood-brain barrier permeability, or cell-to-cell fusion upon viral infection. Here we show the role of MMPs as mediators of virus-induced host neuroinflammatory response in the MHV model. Infection of mice with wild-type MHV-A59 or its isogenic recombinant strains, RSA59 or RSMHV2 significantly upregulated MMP-3, MMP-8, and MMP-14 transcript levels. Functional network assessment with Ingenuity Pathway Analysis revealed a direct involvement of these MMPs in disrupting junctional assembly between endothelial cells via interaction with junctional adhesion molecules and thereby facilitating transmigration of peripheral lymphocytes. Our findings also suggest mRNA upregulation of Park7, which is involved in NADPH oxidase-dependent ROS production, following RSA59 infection. RSA59 infection resulted in elevated mRNA levels of RelA, a subunit of NF-κB. Infection with MHV-A59 is known to generate ROS, and oxidative stress can activate NF-κB. Thus, our findings indicate the existence of a possible nexus between ROS, NF-κB, and MMPs in RSA59-induced neuroinflammation. We also assessed the expression of endogenously produced regulators of MMP activities. Elevated mRNA and protein levels of tissue inhibitors of metalloproteinases 1 (TIMP-1) in MHV-A59 infection are suggestive of a TIMP-1 mediated host antiviral response.

**Importance:** The newly emergent coronavirus has brought the world to a near standstill. In the past, studies have focused on the function of host proteases in virus attachment and entry. Our research indicates the involvement of a group of metal-dependent host proteases in inflammation associated with coronavirus infection. Inflammation is the first response of the host to virus infection. While it helps in restricting the spread and clearance of viral particles, uncontrolled inflammation results in several inflammatory consequences. Therefore, it becomes vital to limit unchecked host immune response. The inhibition of specific metalloproteases represents a potential new therapeutic approach in coronavirus infection and disease outcome.

## 1. Introduction

The newly emergent severe acute respiratory syndrome coronavirus 2 (SARS-CoV-2) (1) and previous encounters with SARS-CoV (2) and middle east respiratory syndrome coronavirus (MERS-CoV) (3) have led to a renewed interest in coronavirus (CoV) research. CoVs are enveloped positive-sense single-stranded RNA virus [ICTV 9^th^ report, 2011; https://talk.ictvonline.org/ictv-reports/]. The first step towards infection of a host requires the virus surface spike (S) glycoprotein (180 KDa) to interact with the host cell surface receptor leading to conformational transitions that promote fusion between viral and host cell membranes (4). The CoV S is composed of two subunits, namely N-terminal S1 (consisting of the receptor-binding domain) and C-terminal S2 (comprising the fusion core domain) subunits. Belonging to the class I viral membrane fusion protein family, CoV S requires cleavage at S1/S2 and S2’ (fusion peptide within S2) sites for effective fusion (4, 5). In general, several cellular proteases such as cell surface serine protease, TMPRSS2 (5-8), intracellular endosomal cathepsins (9-14), or extracellular proteases like trypsin (10, 15, 16) and elastases (16-18) have been detected in priming the S protein for fusion.

Murine hepatitis virus (MHV) is a member of the same beta-coronavirus genus as SARS-CoV-2, MERS-CoV, and SARS-CoV [ICTV 9^th^ report, 2011; https://talk.ictvonline.org/ictv-reports/]. MHV is a widely studied beta-coronavirus model. Being a natural pathogen of mice, MHV damages the liver and central nervous system (CNS), resulting in hepatitis and meningoencephalitis during acute inflammation. MHV-induced chronic inflammation is associated with demyelination concurrent with axonal loss (19). The cleavage of SARS-CoV-2 spike by intracellular proprotein convertase like furin during virus assembly reduces dependence on extracellular proteases for entry (9). Similarly, few MHV S is also cleaved by furin, and furin inhibition prevents cell-to-cell fusion in MHV-A59 strain (20, 21).

Additionally, the role of cellular proteases belonging to the metzincin family called matrix metalloproteinases (MMPs) has been shown in CoV infection (21-23). MMPs are a large family of zinc-dependant endopeptidases, of which 23 members are identified in mouse (24). MMPs are best known for their ability to remodel the extracellular matrix (ECM) proteins. Over the years, the substrate repertoire of MMPs has widened to include cytokines, chemokines, growth factors, and hormones (25). By acting upon cytokines or chemokines immobilized on ECM or cell surface, MMPs release soluble effector molecules for a successful inflammatory response and are involved in autoimmune diseases such as multiple sclerosis, systemic lupus erythematosus, and Alzheimer’s disease (25). Owing to their substrate diversity, MMPs can promote leukocyte migration along the chemotactic gradient by disruption of the blood-brain barrier (25, 26).

Recently, using a broad-spectrum metalloprotease inhibitor, it has been shown that in-vitro infection by MHV strains can be inhibited along with the cell-to-cell fusion without affecting S1/S2 cleavage. The effect on infection varied depending on MHV strains and the cell lines used (21).

Endogenously produced inhibitory proteins called tissue inhibitors of metalloproteinases (TIMPs) regulate the proteolytic activity of MMPs (27). So far, identified four TIMP molecules (TIMP-1, −2, −3, and −4) could inhibit all known MMPs. TIMPs bind reversibly to the catalytic subunit of MMPs in a 1:1 stoichiometric ratio and inhibit their function.

Previous analysis of MMP and TIMP expression in the CNS following infection with neurotropic MHV strain, JHMV showed increased levels of Mmp3, Mmp12, and Timp1 mRNAs, which correlated with high virus replication (23). Moreover, TIMP-1 protein expression was found only in CD4+ T-cells and not CD8+ T-cells, suggesting that TIMP-1 regulates differential CNS distribution of T-cells. TIMP-1, through its inhibition of protease activity, promotes transient accumulation of CD4+ T cells within the perivascular space and controls the trafficking of T cells into the parenchyma (22). Thus, the dynamic regulation of TIMP and MMP levels is an essential factor determining the onset of inflammation. Overall, infection of mice with MHV serves as a useful model to study acute-inflammatory responses and contributions of the immune system in maintaining homeostasis in the inflamed brain.

In this study, we investigated the relationship between the induction of MMP/TIMP expression and inflammation in acute infection of mice with MHV strains. Here we demonstrate that TIMP-1 induction during acute infection following MHV-A59 infection could serve as an antiviral host response to regulate MMP activities. We also determined the role of S protein in inflammation using isogenic recombinant strains RSA59 and RSMHV2 that differ only in the spike gene. Our previous studies have shown that these strains induce similar brain inflammation but differ only in their ability to cause demyelination and axonal loss (28, 29). Here we show that RSA59 and RSMHV2 both induce elevated mRNA expression of MMP-3, MMP-8, and MMP-14. Biological network analysis using Ingenuity Pathway Analysis (IPA) revealed that MMP-3, MMP-8, and MMP-14 aid immune cell infiltration, suggesting a spike-independent role of MMPs in MHV-induced neuroinflammation.

Additionally, RSA59 infection also induced gene transcription of Park7 during acute inflammation. Park7 is involved in the NADPH oxidase-dependant ROS pathway. The transcriptomic analysis also suggests the activation of NF-κB signaling through elevated levels of RelA. RelA is a subunit of NF-κB, and oxidative stress can activate NF-κB. Our study finds a paradigm shift in our understanding of virus-induced inflammatory response by signifying a relationship between RSA59 infection-induced oxidative stress and metalloprotease expression for successful immune cell infiltration.

## 2. Materials and Methods

### Reagents and kits

Gelatin, paraformaldehyde (PFA), and polyvinylidene difluoride (PVDF) membranes were obtained from Merck Millipore, USA. High-capacity cDNA reverse transcription kit (Cat:4368814), DyNAmo ColorFlash SYBR Green qPCR kit (Cat: F-416L), Pierce™ BCA protein assay kit (Cat: 23225), and SuperSignal™ West Pico PLUS Chemiluminescent Substrate (Cat: 34580) were purchased from Thermo Fischer Scientific, USA. EDTA-free protease inhibitor cocktail (Cat:11836170001) was from Roche Diagnostics, Germany. Mouse TIMP-1 antibody (Cat: AF980) was obtained from R&D Systems, USA. Anti-g actin antibody (Cat: BB-AB0025) was purchased from Bio Bharati Life Science, India. Horseradish peroxidase (HRP)-conjugated secondary IgG antibodies, donkey anti-goat (Cat: 705-035-003) and goat anti-rabbit (Cat: 111-035-003) were purchased from Jackson Immuno Research Laboratories, USA. Unless specified, all other reagents were purchased from Sigma while pre-designed primers were ordered from IDT, USA.

### Ethical approval

The use of animals and related experimental techniques were evaluated and sanctioned by the Institutional Animal Care and Use Committee at the Indian Institute of Science Education and Research Kolkata (AUP no. IISERK/IAEC/AP/2017/16). Experiments were performed strictly adhering to the standards of the Committee for the Purpose of Control and Supervision of Experiments on Animals (CPCSEA), India.

### Viruses

A naturally occurring demyelinating strain of MHV, MHV-A59 (Lavi et al., 1984), and two recombinant strains, RSA59 and RSMHV2 (28, 30, 31), were used to infect mice. RSA59 and RSMHV2 are isogenic recombinant strains of MHV-A59 constructed by targeted RNA recombination and differs only in the spike gene as described previously (28, 31). The recombinant demyelinating (DM) RSA59 strain expresses the MHV-A59 spike in the background of the MHV-A59 genome. In contrast, the non-demyelinating (NDM) RSMHV2 strain express MHV-2 [wild-type non-demyelinating MHV (32)] spike protein in the MHV-A59 background. Also, the recombinant strains express enhanced green fluorescence protein (EGFP) for easy in-vitro detection of the virus (30).

### Inoculation of mice

Four-weeks old, MHV-free, male C57BL/6 mice (Jackson Laboratory, USA) were inoculated intracranially with half of 50% lethal dose (LD_50_) of MHV-A59 (2000 PFU), RSA59 (25,000 PFU) or RSMHV2 (100 PFU) respectively and as described previously (28, 31). Three mice (N=3) per group were kept, and 20 μL of the virus prepared in phosphate-buffered saline (PBS) plus 0.075% bovine serum albumin (BSA) was injected into the right cerebral hemisphere of each mouse. Parallelly, we inoculated three mock-infected control mice with only PBS+0.075% BSA. Daily monitoring of mice for disease signs and symptoms was conducted. Control mice were sacrificed between 5-6 days post-infection (p.i). MHV-infected mice were sacrificed at 5-6 (peak of inflammation), 10 and 15 days p.i. For RNA studies, brain and spinal cord tissue samples were flash-frozen in liquid N_2_ and preserved at −80°C until used.

### Estimation of viral replication

The efficacy of virus replication in mice was assessed by routine plaque assay, as described previously (19, 33, 34) with modifications. Briefly, on day 5-6, 10, and 15 p.i, mice were sacrificed, and brain tissues (either half) harvested aseptically in 1mL of isotonic saline containing 0.167% gelatin. Tissues were weighed and stored at −80°C for titer assay. Brain tissues from control and infected mice were homogenized, and 250 μL of serially diluted supernatant was added onto confluent monolayers of DBT cells. Following 75 minutes of viral adsorption, culture media containing 1.4% agarose was overlaid on the monolayers and incubated for 27 hours (MHV-A59/RSA59) or 36 hours (RSMHV2). Cells were fixed with 4% PFA followed by crystal violet staining. Clear regions indicative of plaques were counted, and values [log_10_ pfu/gm tissue] plotted against the time of post-infection. Plaque forming unit or PFU was calculated as the number of plaques times dilution factor (DF) per mL per gram of tissue per mL [PFU= (no. of plaques*DF per ml)/ (tissue weight in gram per ml)].

### Gene expression analysis

Total RNA was extracted from brain tissues of MHV-A59, RSA59 or RSMHV2 infected as well as mock-infected mice using TRIzol reagent (Invitrogen) following the manufacturer’s instructions. RNA concentration was measured using a NanoDrop 2000/2000c Spectrophotometer (Thermo Fisher Scientific), and cDNA was prepared with 1 μg of total RNA using a cDNA reverse transcription kit. Quantitative real-time PCR (RT-qPCR) was performed using SYBR Green dye-based assay in a QuantStudio 3 Real-Time PCR system (Thermo Fisher Scientific) with the following reaction conditions: initial denaturation at 95°C for 7 min, 40 cycles of 95°C for 10 s and 60°C for 30 s, and melting curve analysis at 60°C for 30 s. Reactions were performed in triplicates (n=3). Primer sequences are provided in Table 1. The comparative threshold (ΔΔC_T_) method was used for relative quantification. The mRNA levels of target genes were normalized with the housekeeping GAPDH gene and represented as the relative fold change values compared to their respective mock-infected controls.

**Table 1.**
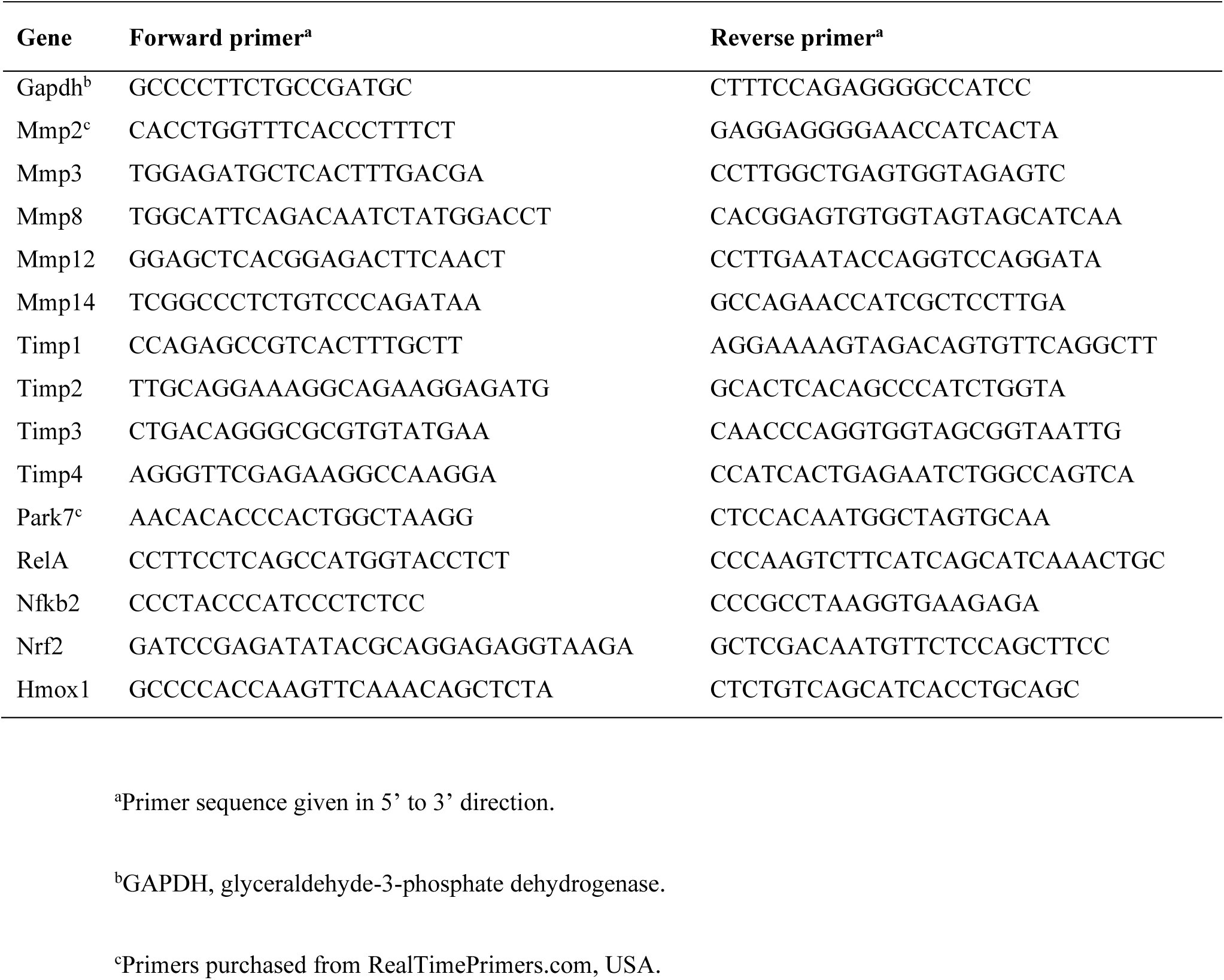
Primer sequence for detection of gene expression by RT-qPCR

### Western blotting

Brain tissues (30 mg) were harvested from mice following transcardial PBS perfusion and flash-frozen in liquid N_2_. Tissues were homogenized (using Qiagen homogenizer) and lysed in 500 uL of RIPA buffer containing protease-cocktail inhibitor and phosphatase inhibitors (1 mM NaVO_4_ and 10 mM NaF) for 1 hr 30 mins with intermittent vortex every 15 mins. The samples were kept on ice during the entire process. Samples were then centrifuged for 15 mins at 13,000 rpm at 4°C to separate the supernatant. The total protein content in the supernatant was estimated with a BCA protein assay kit. For immunoblotting, 60 μg of total protein per sample was resolved by SDS-PAGE on a 12% polyacrylamide gel followed by transfer onto PVDF membranes using transfer buffer (25 mM Tris, 192 mM glycine, and 20% methanol). Membranes were blocked for 1 hr at room temperature in 5% v/v goat serum prepared in TBST (Tris-buffered saline containing 0.1% v/v Tween 20) and subsequently incubated for overnight at 4°C in polyclonal anti-mouse TIMP-1 antibody at 1:1000 dilution in blocking solution. The membranes were washed in TBST and incubated for 1 hr at room temperature with HRP-conjugated donkey anti-goat secondary IgG antibody. As an internal loading control, g-actin was used, and membranes were blocked separately in 5% w/v non-fat skimmed milk in TBST. Polyclonal anti-mouse g-actin antibody (1:5000 dilution) and HRP-conjugated goat anti-rabbit secondary IgG antibody (1:10,000 dilution) were used. The blots were washed in TBST, and the immunoreactive bands visualized using the chemiluminescent HRP substrate. Non-saturated bands were visualized with Syngene G: box Chemidoc system using GENSys Software.

### Affymetrix microarray and pathway analyses

Microarray experiment for mouse gene array was performed with minor modifications as described in detail in our previous studies (35, 36). Briefly, total RNA extracted from spinal cords of RSA59, RSMHV2, and mock-infected mice at day 6 p.i, was subjected to cDNA synthesis using the Ovation Pico WTA System V2 (NuGen Technologies, Inc). Synthesized cDNA was fragmented and biotinylated using FL-Ovation cDNA biotin module V2 (NuGen Technologies, Inc.) followed by hybridization with Affymetrix GeneChip Mouse Gene 1.0 ST Array comprising more than 750,000 unique 25-mer oligonucleotide that contains over 28000 gene-level probe sets (Affymetrix, Santa Clara, CA, USA). Arrays were washed and stained using GeneChip Fluidic Station 450 as per the protocol. Antibody-based amplification of hybridization signals was done using goat IgG (Sigma-Aldrich) and anti-streptavidin biotinylated antibody (Vector Laboratories). Chips were scanned on an Affymetrix Gene Chip Scanner 3000 7G, using Command Console Software. Background correction and normalization was achieved using Iterative Plier 16 with GeneSpring V12.0 software (Agilent Technologies, Inc.). Probe set signals were calculated with the Iterative Plier 16 summarization algorithm, with the baseline to the median of all samples used as the baseline option. Data were filtered by percentile, and a lower cut off was set at 25. A fold change of ≥ 1.5-fold was considered for differential expression of a gene. Statistical analysis using unpaired student t-tests was performed to compare two groups, with p-values ≤ 0.05 considered significant. The list of MMP and TIMP genes from the microarray data was loaded into QIAGEN’s Ingenuity Pathway Analysis software (IPA®, QIAGEN, USA) to perform biological network and functional analyses.

### Statistical analysis

Data shown are mean ± standard error mean (SEM) for all graphs. Unpaired student t-test with Welch’s correction, assuming unequal standard deviations, was performed to examine significant differences between two groups. Multiple comparisons were achieved using ordinary one-way ANOVA, followed by Dunnett’s multiple comparison test. A p-value < 0.05 was considered statistically significant.

### Availability of data

All the data sets used and analyzed in the current study are available from the corresponding author on request.

## 3. Results

### 3.1. Mmp mRNA expression in MHV-A59 infected mice brain

Four weeks old, male C57BL/6 mice inoculated with MHV-A59 (2000 PFU) or mock-infected were sacrificed at day 5-6 (acute), 10 (acute-chronic), and 15 (chronic) post-infection (p.i), and brains were harvested. Routine plaque assay was performed with serially diluted brain homogenates to estimate viral replication. MHV-A59 titer was significant between day 5-6 p.i (Fig. 1, A; p<0.001) and viral particles were below the detection limit at later time points (data not shown). Total RNA was isolated from mock and virus-infected brain tissues for expression analysis of viral nucleocapsid and Mmp genes through RT-qPCR. Primer sequences are given in Table 1. Levels of viral nucleocapsid mRNA (Fig. 1, B; p<0.0001) coincided with viral replication reaching its peak between day 5-6 p.i, which also marks the acute phase of inflammation. MHV-A59 infected mice exhibited neuroinflammation reaching its peak by 5-7 days p.i, which is associated with meningitis, encephalitis, perivascular cuffing, and macrophage/microglia nodule formation (31). We found similar results in paraffin-embedded brain sections stained by hematoxylin-eosin and immunohistochemistry (data not shown). Transcript levels of Mmp2, Mmp3, Mmp8, and Mmp12 were significantly upregulated at day 5-6 p.i (Fig. 1, C-F; p<0.001) and coincided with the peak in viral replication and acute neuroinflammation. Additionally, Mmp14 mRNA was significantly upregulated only at later time points, between days 10-15 p.i (Fig. 1, G; p<0.001). Collectively, MHV-A59 infection-induced the upregulation of Mmp2, Mmp3, Mmp8, Mmp12, and Mmp14.

**Fig. 1.**
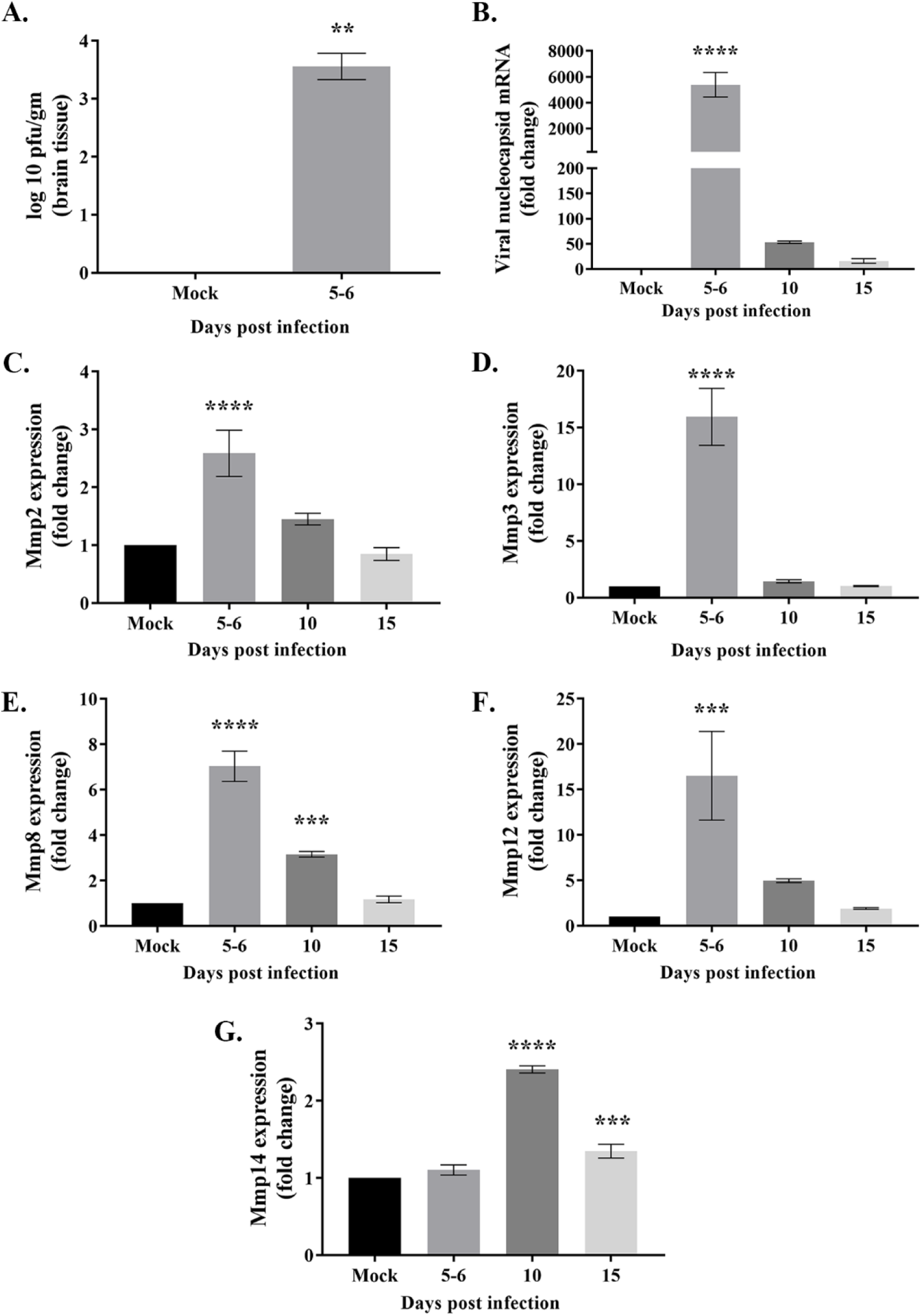
Upregulation of Mmp2, Mmp3, Mmp8, Mmp12, and Mmp14 mRNAs in the brain of MHV-A59 infected mice. Brain tissues from mock- and MHV-A59 (2000 PFU)-infected mice at different days post-infection (p.i) were harvested, and viral replication was estimated by routine plaque assay. RNA isolated from brain tissues was subjected to cDNA synthesis. An equal amount of cDNA template was used for RT-qPCR. Gene expression was normalized to GAPDH and fold-change values obtained using ΔΔC_t_ method. A: MHV-A59 titer peaked between day 5-6 p.i. B: Viral nucleocapsid mRNA levels peaked at day 5-6 p.i like viral replication. C-F: Mmp2, Mmp3, Mmp8, and Mmp12 mRNA levels elevated between 5-6 days p.i. and coincided with viral replication peak. G: Membrane-associated MMP-14 mRNA levels peaked only at later stages p.i. Data shown are mean ± SEM from two independent biological experiments with nine technical replicates. A significant difference between the two groups was compared with the student t-test. Multiple group comparison was made with ordinary one-way ANOVA followed by Dunnet’s test. A p-value of < 0.05 was considered statistically significant (**, p<0.01; ***, p<0.001; ****, p<0.0001).

### 3.2. Induction of TIMP-1 expression following MHV-A59 infection

Tissue inhibitors of metalloproteinases or TIMPs are endogenous protein regulators of MMPs. To understand the regulation of MMPs upon MHV-A59 infection, we also considered the gene expression of TIMPs. As described above, total RNA from brain samples of mock and MHV-A59 infected mice were subjected to RT-qPCR using specific primers (Table 1) to determine the transcript levels of Timp1, Timp2, Timp3, and Timp4. MHV-A59 infection resulted in significant upregulation of Timp1 mRNA at day 5-6 p.i (Fig. 2, A; p<0.001), while mRNA levels of Timp2, Timp3, and Timp4 remained significantly downregulated (Fig. 2, C-E; p-values varies as <0.05 to <0.0001). While Timp1 mRNA followed a similar expression pattern as the Mmps following MHV-A59 infection-induced inflammation, its protein levels remained high throughout post-infection, as shown in the representative figure (Fig. 2, B). Overall, MHV-A59 resulted in elevated TIMP-1 levels in the brain of infected mice.

**Fig. 2.**
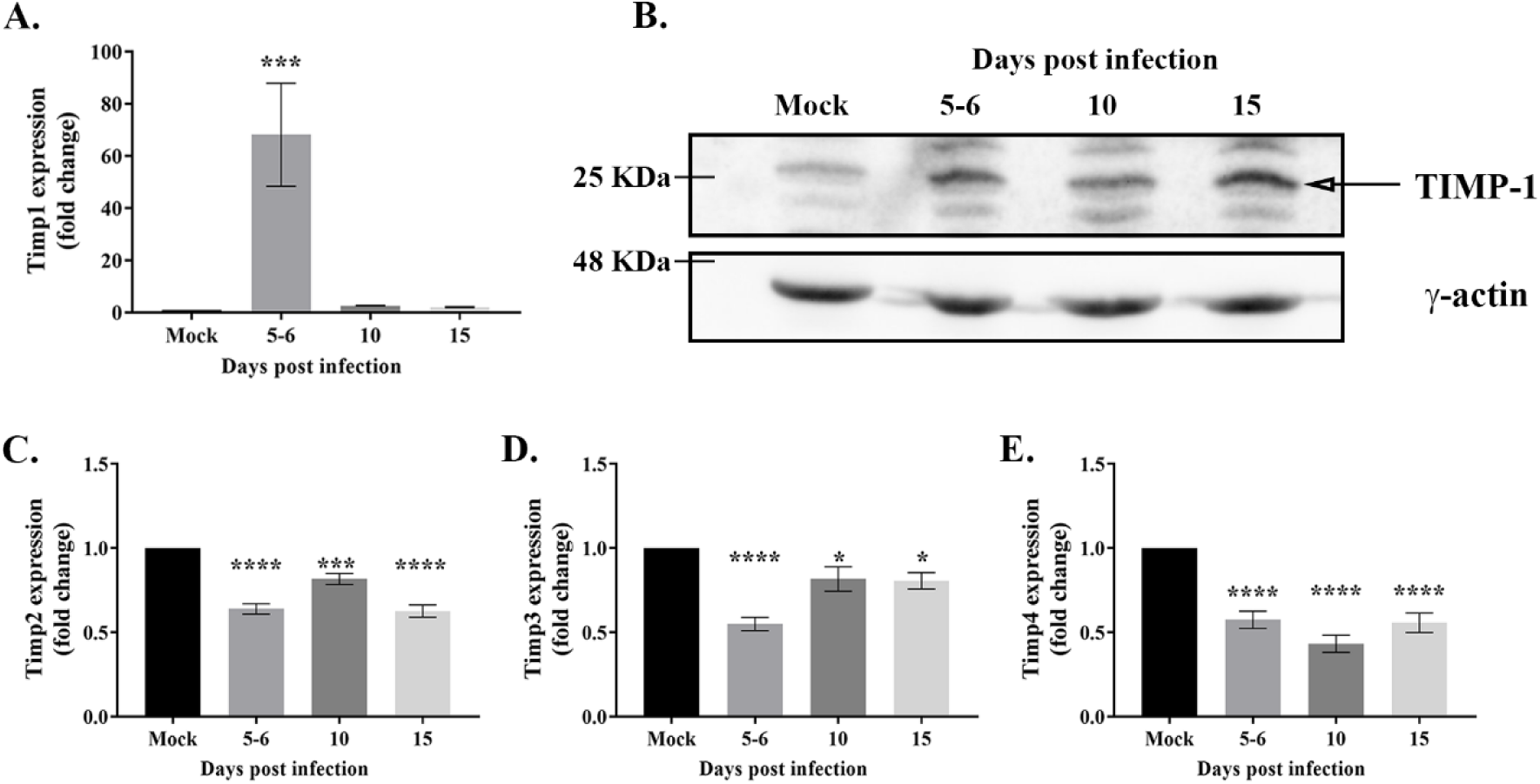
Preferential upregulation of TIMP-1 following MHV-A59 infection. Data analysis of RT-qPCR revealed a significant upregulation in Timp-1 mRNA levels at day 5-6 p.i in MHV-A59 compared with mock-infected samples (A). Representative immunoblot assay from two independent experiments showed elevated protein levels of TIMP-1 at all examined days following MHV-A59 infection (B). In contrast, Timp-2, −3, and −4 mRNAs remained downregulated throughout p.i (C-E). Graphs show mean ± SEM values from two independent experiments with nine technical replicates. Ordinary one-way ANOVA followed by Dunnet’s test was performed for multiple group comparisons. Statistical significance was considered for p values < 0.05 (*, p<0.05; ***, p<0.001; ****, p<0.0001).

### 3.3. Similarities in the induction of MMP and TIMP expression following RSA59 and RSMHV2 infection

To determine whether the spike (S) protein has any role in inducing Mmp and Timp expression, we employed isogenic recombinant strains RSA59 and RSMHV2 that differ in their spike protein. In our previous publications (35, 36), using Affymetrix microarray analysis with RNA isolated from spinal cords (day 6 p.i), we have shown that both the recombinant strains induced the upregulation genes related to immune modulation during acute infection. Using the same Affymetrix microarray data, we analyzed Mmp and Timp expression levels in RSA59 and RSMHV2 infection. A threshold of ≥1.5 fold-change with p-values ≤0.05 was considered significant for determining differential gene expression. The analysis revealed that in both strains, there were no significant differences in the expression of Mmp and Timp genes compared with mock samples during the acute infection in the spinal cord. To validate the findings from microarray data, we performed RT-qPCR from brain samples of RSA59, RSMHV2, and mock-infected mice sacrificed at day 5-6, 10, and 15 p.i. Brain samples were also harvested for titer assay to estimate viral replication. Like the parental MHV-A59 strain, viral titer and nucleocapsid mRNA levels peaked by day 5-6 p.i in both the recombinant strains as demonstrated in the representative graphs (Fig. 3, A-C). Although we detected no nucleocapsid mRNAs between 10-15 days p.i in RSMHV2, its presence was observed at day 10 p.i in RSA59 (data not shown). This data corroborates with previous findings that demyelinating RSA59 persists in the brain while RSMHV2 does not persist or, if present, nucleocapsid level is significantly low compared with RSA59 (28). Also, both the strains showed similar kinetics in Mmp gene expression like their parental MHV-A59. Results from RT-qPCR demonstrated significant upregulation of Mmp3, Mmp8, and Mmp14 mRNA levels in both the strains when compared with mock-infection between day 5-6 p.i (Fig. 4, A-F; p-value varies as <0.05 to <0.0001). Additionally, the two strains did not show any difference in altering the levels of Timp1 (Fig. 4, G & H; p<0.0001). Both RSA59 and RSMHV2 do not differ significantly in their ability to induce inflammation in the brain (29, 37). Therefore, we performed biological pathway analysis to understand the involvement of MMP-3, −8, and −14 in the onset of virus-induced neuroinflammation.

**Fig. 3.**
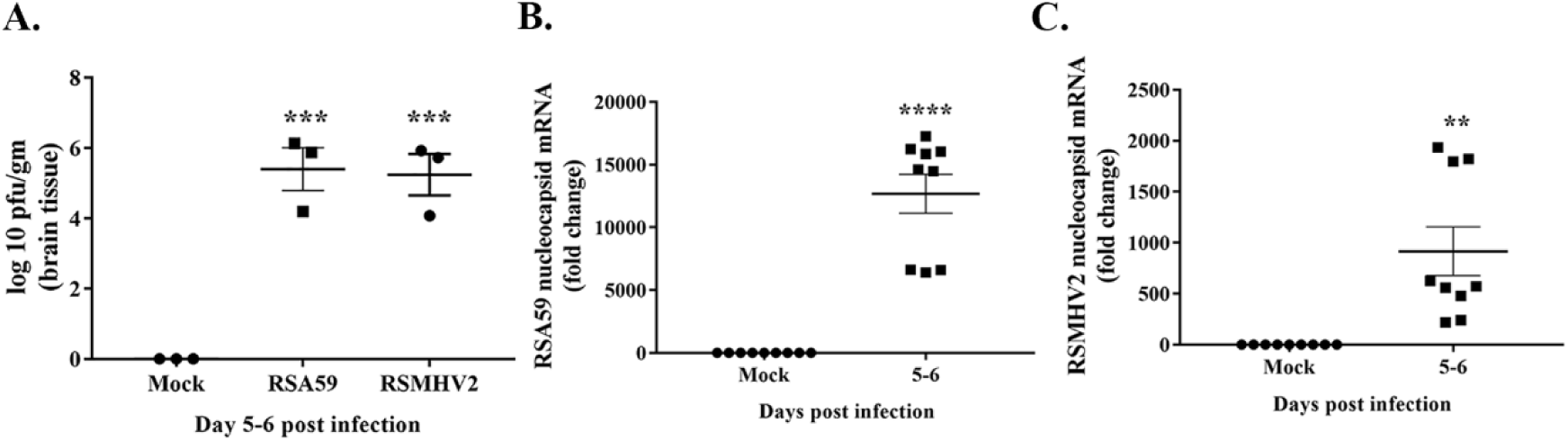
Viral replication and nucleocapsid mRNA expression in isogenic recombinant MHV strain at 5-6 days post-infection. RSA59 and RSMHV2 are isogenic recombinant strains of the wild-type MHV-A59 and differs only in the spike gene. Brain samples from mice infected with RSA59 (25000 PFU) or RSMHV2 (100 PFU) were harvested at different days p.i for routine plaque assay and total RNA extraction. A: Both RSA59 and RSMHV2 showed peak viral replication between day 5-6 p.i. No detectable viral particles were observed at later time points in a routine plaque assay (data not shown). RT-qPCR data showed significantly elevated mRNA levels of viral nucleocapsid gene in both RSA59 (B) and RSMHV2 (C) at early days p.i, coinciding with peak levels of viral particles as detected in a plaque assay. Data shown are mean ± SEM from three independent experiments having three technical replicates each. Multiple group comparison was made with ordinary one-way ANOVA followed by Dunnet’s test. A p-value of < 0.05 was considered statistically significant (***, p<0.001; ****, p<0.0001).

**Fig. 4.**
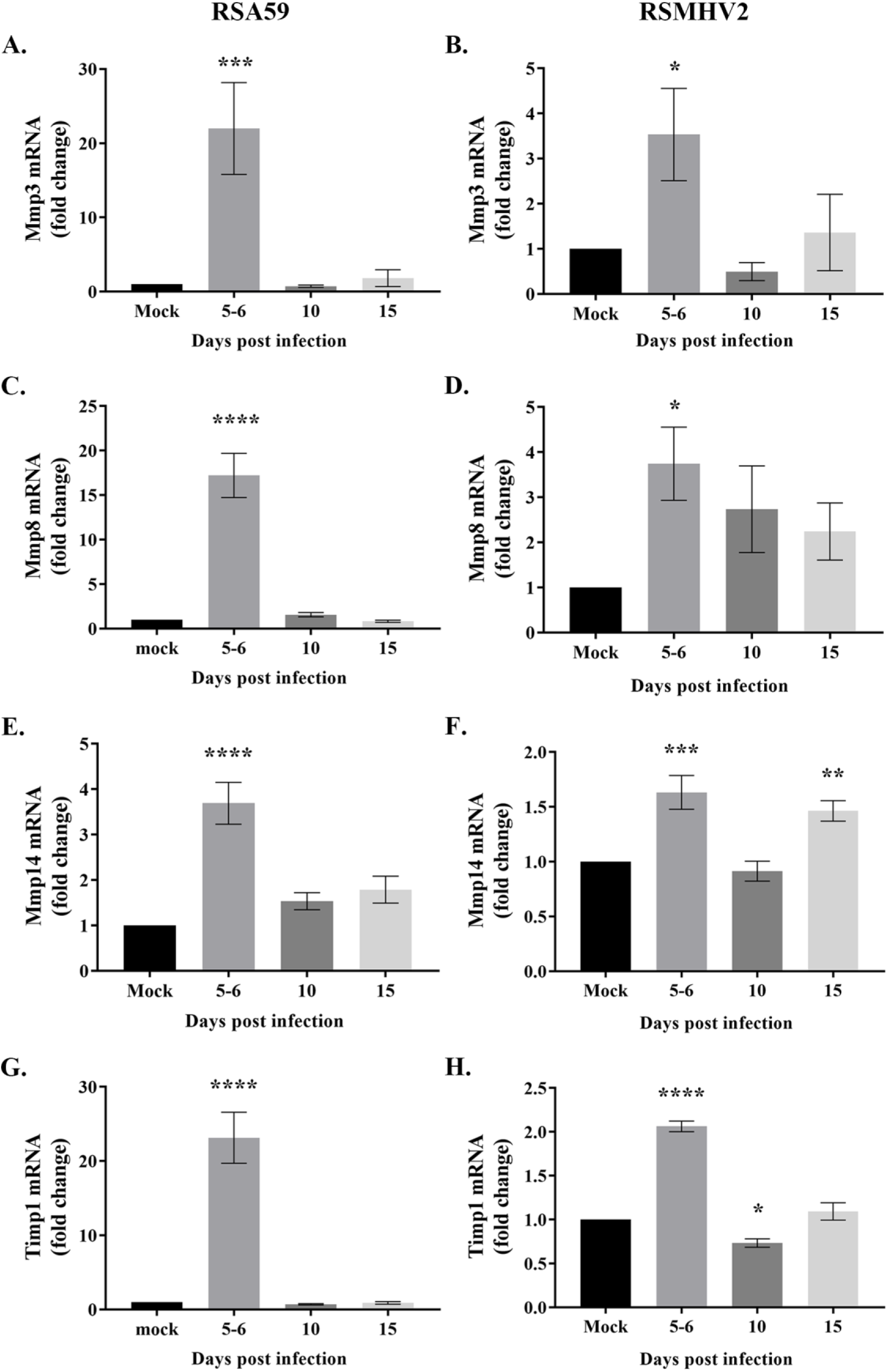
Elevated brain levels of Mmp3, Mmp8, Mmp14, and Timp1 mRNAs at the peak of inflammation following infection with isogenic recombinant strains, RSA59 and RSMHV2. Like their parental wild-type MHV-A59 strain, both RSA59 and RSMHV2 induced similar expression patterns of Mmp and Timp genes, as evident from RT-qPCR. A-F: Mmp3, Mmp8, and Mmp14 transcript levels were significantly upregulated between day 5-6 p.i, coinciding with the peak in the inflammatory response. G-H: Additionally, Timp1 mRNA was also upregulated in both the strains and followed a similar expression trend as Mmps. Data shown are mean ± SEM from three (A-F) and two (G-H) independent experiments having three technical replicates each. Ordinary one-way ANOVA followed by Dunnet’s test was performed for multiple group comparisons. Statistical differences at p < 0.05 was considered significant (*, p<0.05; **, p<0.01; ***, p<0.001; ****, p<0.0001).

### 3.4. Classical pathways representing leukocyte adhesion and diapedesis activated in RSA59 and RSMHV2 infection

Biological and functional network analysis were performed for Mmp3, Mmp8, and Mmp14 genes using QIAGEN’s Ingenuity Pathway Analysis (IPA) software. IPA analysis identified that these MMPs could influence several canonical pathways associated with an immune response such as leukocyte extravasation signaling, granulocyte and agranulocyte adhesion and diapedesis (Fig. 5, A). Also, the top disease pathways involved both inflammatory response and immune cell trafficking (Fig. 5, B). Furthermore, IPA revealed that MMPs facilitate the transmigration of firmly adhered granulocytes (Fig. 6, A) and agranulocytes (Fig. 6, B) across the endothelial cells in the blood vessel. Results indicated a relationship between MMPs and junctional proteins like claudin, cadherin, and platelet-derived endothelial cell-adhesion molecule. Collectively, IPA analysis suggests the role of MMP-3, MMP-8, and MMP-14 in peripheral immune cell migration during MHV infection.

**Fig. 5.**
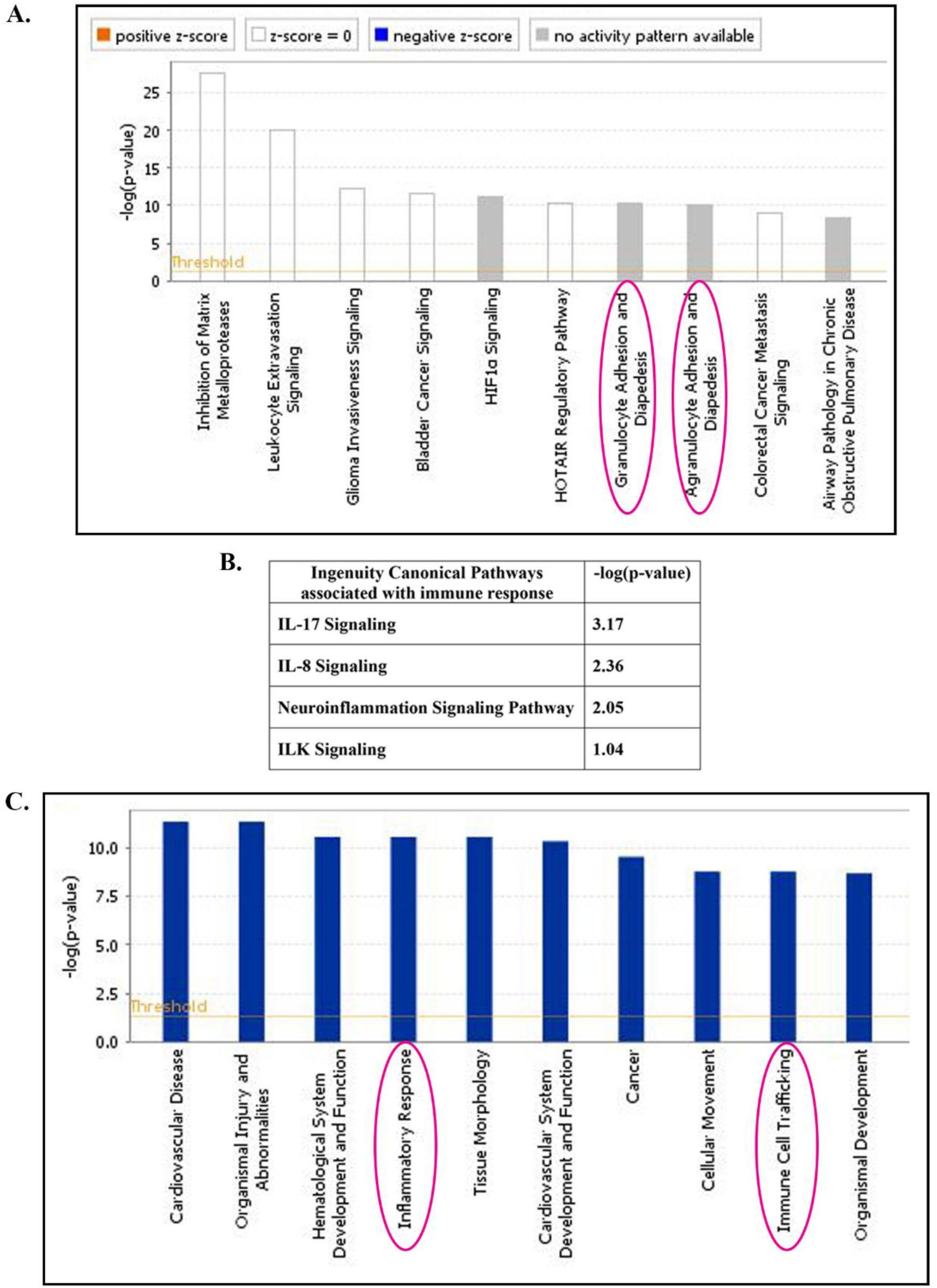
Analysis of significantly upregulated Mmp genes in RSA59 and RSMHV2 infection compared to mock-infected mice. As shown in Fig. 4, transcript levels of Mmp3, Mmp8, and Mmp14 genes were elevated during the acute stage of infection in both RSA59 and RSMHV2 infected mice. These genes were further analyzed by Ingenuity Pathway Analysis (IPA) software and found to be involved in several canonical and disease pathways. A: Among the top canonical pathways, the involvement of Mmp3, Mmp8, and Mmp14 genes were observed in granulocyte and agranulocyte adhesion and diapedesis. B: Additionally, other inflammation-related canonical pathways depicted in IPA that involves Mmps are IL-17, IL-8, ILK-signaling, or neuroinflammation signaling. C: Moreover, the role of Mmp3, Mmp8, and Mmp14 in the inflammatory response and immune cell trafficking featured among the top disease pathways upon viral infection. A smaller p-value means that the association between the analyzed genes in MHV infection with the known set of genes implicated in that given pathway is significant and less likely to be random. P-values <0.05 are considered statistically significant, and the threshold line represents the p-value is 0.05. Pathways are ranked according to their p-value and shown as bars. Given are only the top ten pathways for canonical and disease analysis graphs (A and C).

**Fig. 6.**
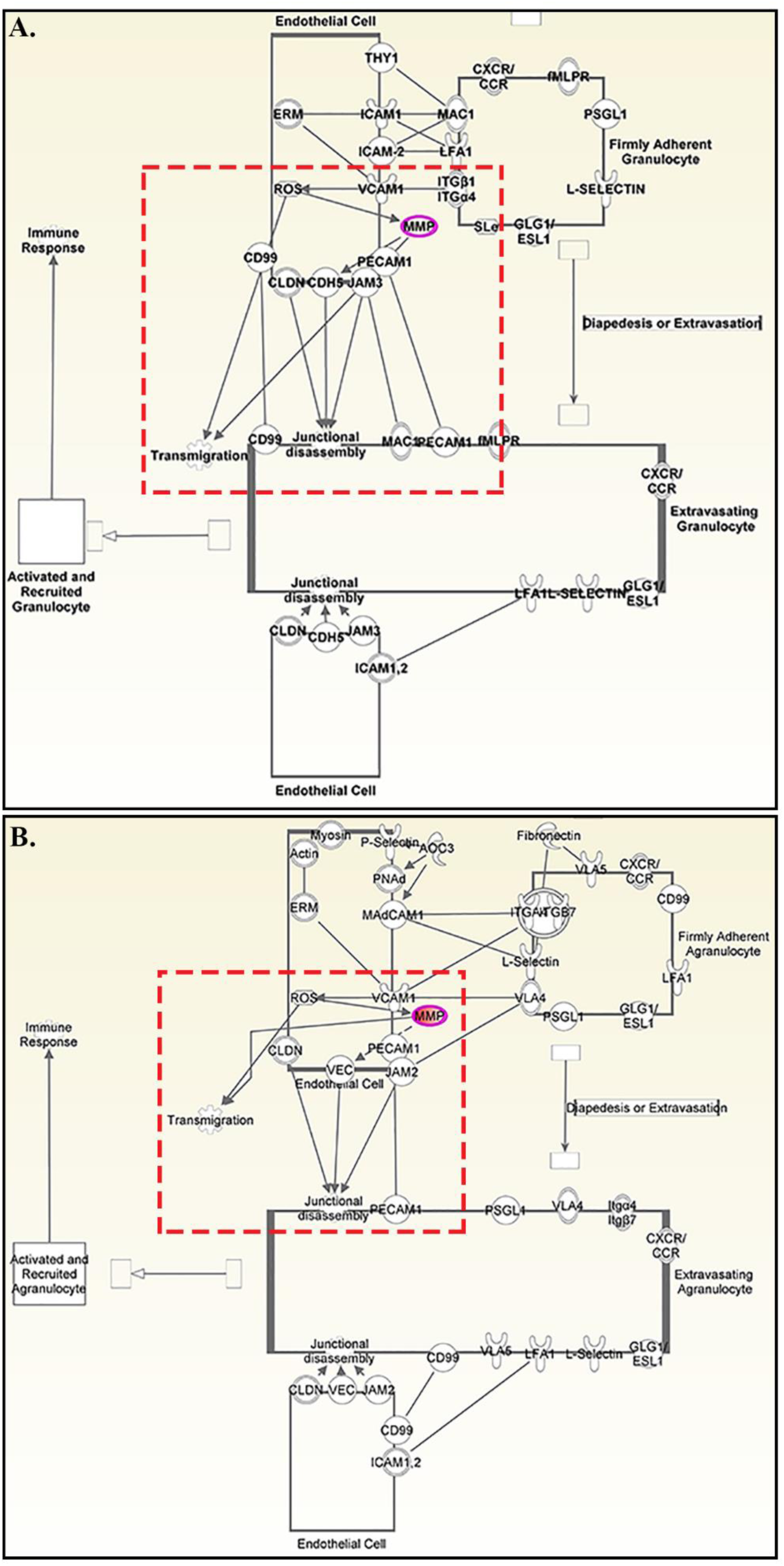
Network analysis of Mmp3, Mmp8, and Mmp14 in RSA59 and RSMHV2 infection. The path designer function of the IPA software was used to dissect further the direct and indirect relationships that exist between these Mmp genes and inflammatory response with a focus on immune cell trafficking. Results demonstrate that Mmps facilitate the transmigration of granulocytes (A) and agranulocytes (B) firmly adhered to endothelial cells in the blood vessel. As observed during granulocyte extravasation (A), MMPs mediate junctional disassembly between endothelial cells via interacting with adhesion molecules such as claudin (CLDN), cadherin 5 (CDH5), endothelial cell adhesion molecule (PECAM1), and junctional adhesion molecules (JAM). Nodes in the network represent the genes (or their corresponding proteins), and the lines connecting the nodes indicate the kind of interaction between the genes (direct is the solid line; indirect is a dotted line). Red dotted rectangles highlight the interaction between MMPs and different molecules.

### 3.5. RSA59 infection induced increased transcription of oxidative and anti-oxidative pathway genes

Brain samples from mice infected with RSA59 and sacrificed at day 5-6 and 10 p.i, were harvested for total RNA isolation followed by cDNA synthesis. Mock-infected samples were kept in parallel. We performed RT-qPCR using primers (Table 1) specific for genes involved in the oxidative and anti-oxidative pathways. Transcript levels of Parkinson’s disease 7 (Park7) gene were significantly upregulated following RSA59 infection and remained elevated p.i compared to mock-infected samples (Fig. 7, A; p<0.05). RelA, a subunit of NF-κB, also showed elevated mRNA levels during the acute infection, i.e., 5-6 days p.i (Fig. 7, B; p<0.0001). On the contrary, mRNA levels of Nfκb2, a negative regulatory subunit of NF-κB, remained unchanged p.i (Fig. 7, C). Similar to another study of our lab (unpublished data), we detected significantly high mRNA levels of nuclear factor erythroid 2-related factor 2 (Nrf2) and heme oxygenase-1 (Hmox1) genes during the acute disease phase (Fig. 7, D-E; p<0.0001). Overall, RSA59 infection-induced simultaneous transcription of genes such as Park7 and Nfκb that are involved in the oxidative pathway as well as anti-oxidative pathway genes like Nrf2 and Hmox1.

**Fig. 7.**
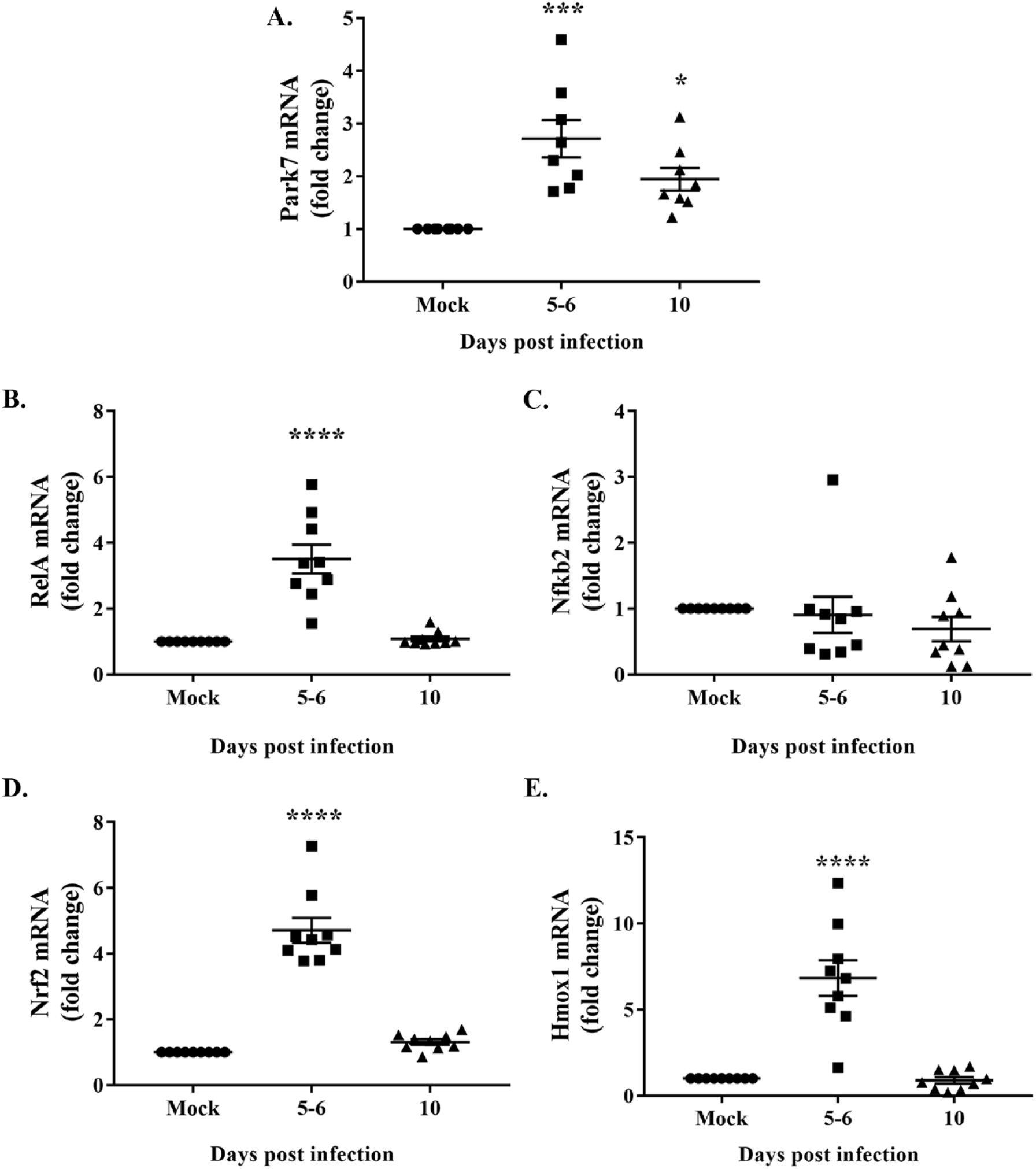
Upregulation of gene transcripts involved in oxidative (A-C) and anti-oxidative (D & E) pathways during RSA59 infection. Total RNA isolated from brain samples of mice infected with RSA59 (25000 PFU) or mock-infected was subjected to cDNA synthesis, and subsequently, RT-qPCR was performed. A: Data analysis revealed elevated mRNA levels of Park7 during acute (5-6 days p.i) and acute-chronic (day 10 p.i) disease phase. B: mRNA upregulation was detected for RelA, a subunit of the NF-κB transcription factor. C: In contrast, no change was observed in the mRNA level of Nfκb2, a negative regulator of NF-κB. D & E: Moreover, RSA59 infection also induced increased transcription of Nrf2 and Hmox1 genes. Data shown are mean ± SEM from two independent experiments. A significant difference between multiple groups was compared with ordinary one-way ANOVA, followed by Dunnet’s test. A p-value of < 0.05 was considered statistically significant (*, p<0.05; ***, p<0.001; ****, p<0.0001).

## 4. Discussion

The current study investigates the regulation of MMPs and their inhibitors during CNS infection with MHV strains. Wild-type MHV-A59 and its isogenic recombinant strains, RSA59 and RSMHV2, were assessed to determine their regulation of MMP and TIMP mRNA expression. Kinetics of MMP and TIMP expression, mostly upregulated during the acute disease phase, was similar in both the parental MHV-A59 and recombinant strains. Here, we show that upregulation of Mmp3, Mmp8, and Mmp14 mRNAs during MHV infection irrespective of differences in the surface spike protein. MMP-3 (stromelysin) is produced by leukocytes and endothelial cells and can act on multiple collagen substrates. Both neutrophils and macrophages produce MMP-8 (neutrophil collagenase), while MMP-14, which is membrane-associated, promotes the conversion of inactive MMP-2 to its active form. Both MMP-8 and MMP-14 can cleave multiple collagen substrates (27). Collectively, the ability of MMP-3, −8, and −14 to modulate the ECM is indicative of their involvement in the migration of peripheral immune cells. These we further confirmed from biological and functional network evaluation using Ingenuity Pathway Analysis (IPA). IPA analysis suggests a direct role of MMP-3,-8, and −14 in granulocyte and agranulocyte transmigration across the endothelial cells by disrupting claudin, cadherin, or junctional adhesion molecules mediated tight junctional assembly. Moreover, it explains the similarity observed in CNS inflammation with both the recombinant strains RSA59 and RSMHV2 despite having differences in their S proteins. Past studies associating overproduction of MMPs with increased blood-brain-barrier permeability and thereby, enhanced inflammation of neurotropic viruses (38, 39), further supports our observation.

In previous studies involving JHMV strains, it has been shown that MMP-3, MMP-12, and TIMP-1 mRNA expression correlates with viral virulence and viral load (23). Additionally, the role of TIMP-1 in inhibiting protease activity and thereby controlling the differential infiltration of T-cell subsets during virus encephalitis was also demonstrated (22). We found Timp1 significantly upregulated in both the strains like the parental MHV-A59. Our study adds further knowledge in understanding the role of TIMP-1. The present work suggests that Timp1 upregulation could be part of a classical host defense mechanism against virus-induced upregulation of different metalloproteases. Whether the downregulation of Timp2, Timp3, and Timp4 is purposely mediated by MHV or there exists a feedback mechanism to maintain a balance among different TIMPs is an exciting aspect to address. Moreover, our study further sheds light into the viral-induced oxidative stress pathway and neuroinflammation.

Oxidative stress is an intrinsic mode of tissue damage and also occurs in MHV-A59 infection (40). A NAD-dependent deacetylase, SIRT1 involved in cell stress responses, prevents neuronal injury, decrease oxidative stress, and induce mitochondrial function-associated proteins upon MHV-A59 infection. Oral treatment of MHV-A59 infected mice with a SIRT1 activating compound increased expression of mitochondrial enzymes such as succinate dehydrogenase (SDH) and superoxide dismutase 2 (SOD2) and significantly reduced ROS levels. In our current study, RSA59 infection increased transcript levels of Park7, which is involved in oxidative stress. Park7 has a double-sword effect. In lower ROS concentration, it can affect NADPH oxidase by phosphorylating its p47^phox^ subunit during NADPH oxidase activation, which is crucial for NADPH oxidase-dependant ROS production (41). In one of our studies (unpublished data), we show that Park7 also induces the anti-oxidative pathway via Nrf2 and Hmox1activation during higher cellular ROS concentration.

Activation of metalloproteases via oxidative pathways has been demonstrated in the past (42-44). ROS-induced oxidative stress can also activate NF-κB signaling (45). The nuclear factor kappa-light-chain-enhancer of activated B cells (NF-κB) acts as a transcription factor and is known to induce inflammation-related genes. RelA, a subunit of NF-κB, which gets activated in the canonical pathway via toll-like receptors that recognize pathogenic patterns (46), showed increased mRNA levels following RSA59 infection. On the other hand, Nfκb2, which acts as both a precursor and suppressor of NF-κB (46), demonstrated unchanged mRNA levels upon infection. Previously, it has been documented that NF-κB can induce MMP genes (47, 48). Therefore, our result indicates that Park7 mediated ROS generation leads to the induction of MMP genes via NF-κB signaling during MHV-induced acute disease (Fig. 8).

**Fig. 8.**
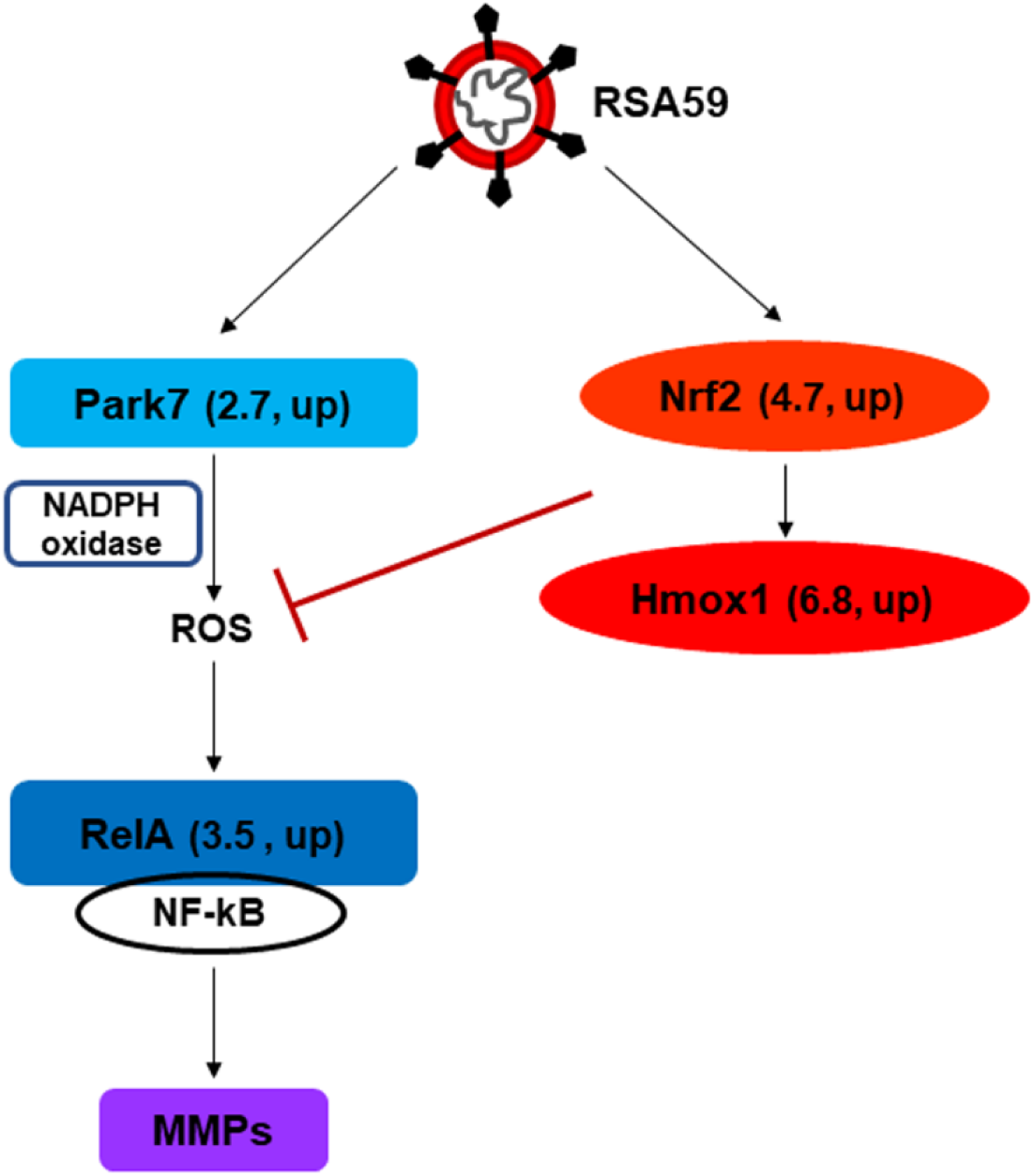
Graphical abstract depicts the relationship between ROS, NF-κB, and induction of MMP genes following RSA59 infection. RSA59 induced the upregulation of Park7 (Parkinson’s disease 7) in the brain of virus-infected mice compared with mock samples. Park7 is known to activate NADPH oxidase-dependant ROS production. In the downstream of the ROS pathway lies nuclear factor kappa B or NF-κB. RSA59 infection also upregulated the transcripts of RelA, a subunit of NF-κB. The transcription factor NF-κB can induce many inflammation-related genes, including MMPs. Additionally, RSA59 infection increased mRNAs of nuclear factor erythroid 2-related factor 2 (Nrf2) and its dependant heme oxygenase-1 (Hmox1) genes that are involved in the anti-oxidative pathway. The numbers indicate mRNA fold-change along with its regulation (up or down) as detected in RT-qPCR in Fig. 7.

We also found that RSA59 infection-induced upregulation of Nrf2 and Hmox1 genes. The anti-oxidative pathway mediated by nuclear factor erythroid 2-related factor 2 (Nrf2) and its dependant heme oxygenase-1 (Hmox1) (49), could therefore play an essential role in restoring homeostasis through inhibition of ROS overproduction. One limitation of this study that will be addressed in our future experiments is that the interplay between ROS and MMPs has not been validated using inhibitors of ROS as positive controls.

In previous studies (50-52) involving MHV-A59, it has been shown that virus infection reduced expression of connexins (Cxs) that form intercellular gap junctional channels and thereby disrupt functional communications between CNS glial cells and fibroblasts. However, the mechanism through which Cx trafficking is altered is not well understood. The inhibition of MMPs is shown to increase Cx levels in a hypoxia model (47). Our future experiments are directed towards understanding the interaction between MMPs and Cxs to explore the underlying mechanism of MHV-induced alteration of gap junctional communication.

While the dependence of beta-coronaviruses on cellular proteases for successful entry is widely studied, our findings demonstrate an essential role of cellular proteases in mediating inflammation. Our data suggest that MMP-3, MMP-8, and MMP-14 might play prominent roles in MHV-induced neuroinflammation. Another interesting finding is the existence of a possible nexus between ROS, NF-κB, and MMPs during MHV infection. Further studies needed to enhance our understanding of the diverse role played by MMPs in inflammation and in developing strategies to interrupt this process, thereby influencing the outcome of CNS inflammatory diseases. Again, in the human beta-coronavirus SARS-CoV-2, which causes acute respiratory illness, it will be fascinating to explore the role of MMPs or metalloproteases in general.

## Author Contributions

Conceptualization, designing, and planning of all experiments conducted by S.S. and J.D.S; S.S performed all the animal experiments; S.A performed the microarray experiments, network construction using IPA and provided help to S.S and J.D.S for data interpretation; D.B helped with the RT-qPCR experiments and its data analysis; S.S and J.D.S participated in data analysis and data interpretation; S.S and J.D.S drafted the manuscript; J.D.S critically reviewed the manuscript and supervised all aspects of this work. All authors have read and agreed to the submission of this manuscript.

## Acknowledgments

We thank the Ministry of Education, India, and the Department of Science & Technology for fellowships to S.S and D.B, respectively. We also thank the Department of Biological Sciences, IISER-K, for providing the necessary laboratory facilities. We acknowledge the support of the Cancer Genomics Center, Thomas Jefferson University, Philadelphia, USA, for the Gene Chip Microarray facility. The authors thank the IISER-K Animal Facility. Authors also thank the IISER-K SyMeC Lab for providing access to its RT-qPCR facilities.

